# Comprehensive analysis of RNA-seq kits for standard, low and ultra-low quantity samples

**DOI:** 10.1101/524439

**Authors:** Marie-Ange Palomares, Cyril Dalmasso, Eric Bonnet, Céline Derbois, Solène Brohard-Julien, Christophe Ambroise, Christophe Battail, Jean-François Deleuze, Robert Olaso

## Abstract

High-throughput RNA-sequencing has become the gold standard method for whole-transcriptome gene expression analysis, and is widely used in numerous applications to study cell and tissue transcriptomes. It is also being increasingly used in a number of clinical applications, including expression profiling for diagnostics and alternative transcript detection. However, despite its many advantages, RNA sequencing can be challenging in some situations, for instance in cases of low input amounts or degraded RNA samples. Several protocols have been proposed to overcome these challenges, and many are available as commercial kits. In this study, we comprehensively test three recent commercial technologies for RNA-seq library preparation (TruSeq, SMARTer and SMARTer Ultra-Low) on human reference tissue preparations, using standard (1*μ*g), low (100 and 10 ng) and ultra-low (< 1 ng) input amounts, and for mRNA and total RNA, stranded or unstranded. The results are analyzed using read quality and alignment metrics, gene detection and differential gene expression metrics. Overall, we show that the TruSeq kit performs well with an input amount of 100 ng, while the SMARTer kit shows degraded performance for inputs of 100 and 10 ng, and the SMARTer Ultra-Low kit performs relatively well for input amounts < 1 ng. All the results are discussed in detail, and we provide guidelines for biologists for the selection of a RNA-seq library preparation kit.

## Introduction

RNA molecules play an essential role in numerous biological processes. With the recent evolution of next-generation sequencing technologies, it is now easier than ever before to investigate the large diversity of RNA species using high-throughput RNA sequencing (RNA-seq). Unlike older techniques such as quantitative reverse transcription and microarrays, RNA-seq is an open platform in the sense that it does not depend on genome annotation or predefined and species specific probes for transcript measurement, thus allowing the detection of both known and novel transcripts, including variants and rare transcripts^1^. RNA-seq has been shown to have a greater dynamic range than microarrays, increasing the potential for the detection of variation within samples^2^. This technology can identify tens of thousands of differentially expressed genes and their isoforms. It can also detect mutations and germline variations for hundreds of thousands of expressed genetic variants, and can detect gene fusions and splice variants^3, 4^. In addition, RNA-seq can identify various classes of non-coding RNAs, such as microRNAs, PIWI-interacting RNAs, and tRNAs^4^. RNA-seq techniques are being increasingly used in clinical applications. For instance, the recent breast cancer guidelines support the usage of mRNA based prognostic assays to assist in treatment decisions, alongside other clinico-pathological factors^5^. These assays could also provide a better view of alternative transcripts variants which arise from splicing alterations or structural variants, and are implicated in a range of human diseases such as developmental disorders^6^, neurodegenerative disorders^7^ and cancers^8^. Thus, RNA-seq will most likely transition in the very near future from a discovery tool to diagnostic tool with clinical applications such as patient stratification, diagnosis and personalized treatment^1^. Producing high quality RNA-seq libraries and sequencing results can be challenging due to the multiple factors and conditions involved. For instance, in many cases it is necessary to reduce the representation of abundant ribosomal RNAs (rRNA) in libraries prior to sequencing. Sometimes libraries have to be prepared from poor quality samples such as formalin-fixed, paraffin-embedded (FFPE) samples. In other cases, working with sorted cell subtypes, minute tissue samples or even single cells can result in low or ultra low sample input amounts. To overcome these challenges, several methods and kits have been developed, each with its own strengths and weaknesses. The relative merits of each method compared to a standard high-quality and high quantity input should be determined by a careful comparison of multiple metrics^9–12^. Several studies have investigated the impact of degraded RNA input^13–17^, while others have focused on low RNA inputs^16, 18–20^, with some studies focusing on both aspects^16, 17^, or on the general properties of the RNA-seq protocols^3^. Here, in the light of recent developments and progress in RNA-seq library protocols, we use human input samples to evaluate three different recently developed commercial kits used for RNA-seq library preparation: TruSeq (Illumina), SMARTer (Clontech/Takara Bio) and SMARTer Ultra-Low (Clontech/Takara Bio). Furthermore, we test these kits for unstranded and stranded conditions, mRNA and total RNA input selection and for three different quantities of input material: standard (1 *μ*g), low (100 ng and 10 ng) and ultra-low (< 1 ng). The results are then analyzed using read quality and alignments metrics, gene detection and differential gene expression metrics. In total, we sequence 80 different libraries representing the different preparation kits / input material combinations using high-throughput sequencing. Based on the data generated and the gene detection and differential expression analysis, we discuss the performance of the different kits and input amounts, and their impact on investigating the transcriptomic landscape in general.

## Results

### RNA samples, sequencing protocols and study design

In order to evaluate the performance of RNA-seq methods for profiling normal, low and ultra-low quantity samples, we compared the quality metrics, gene detection and differential analysis capacities of three different groups of RNA extraction kits. The different conditions used are detailed in Table 1. The first group is based on the widely used Illumina TruSeq technology which we used for total RNA and mRNA extraction, stranded or unstranded, with input amounts of 1*μ*g and 100 ng. The second group is based on SMARTer technology which we used for total RNA extraction, performing the ribosomal RNA depletion with two different kits (RiboGone and RiboZero) and using input amounts of 100 ng and 10ng. The third group is based on a combination of SMARTer Ultra-Low plus Nextera technologies for ultra-low input amounts of 750 and 130 pg. For the RNA input material, we used two human RNA sample preparations previously used in reference RNA-seq studies^9, 11, 12^ (see methods). All samples were sequenced in duplicate giving a total of 80 sequenced libraries. Paired-end reads were then checked for quality, aligned to the human genome and counts were generated for all human genes. Lastly, for the gene detection and differential analysis studies, mapped reads were sampled at four different levels.

**Table 1.** Overview of the different RNA preparation kits and conditions analyzed in this study.

| Kit | Manufacturer | RNA type | Input (ng) | Stranded | Acronym |
| --- | --- | --- | --- | --- | --- |
| **Illumina technology** |  |  |  |  |  |
| mRNA TruSeq | Illumina | mRNA | 1000 | no | mRNATruseq_1ug |
| Stranded mRNA TruSeq | Illumina | mRNA | 1000 | yes | ssmRNATruseq_1ug |
| Stranded total RNA TruSeq | Illumina | total RNA | 1000 | yes | ssTotRNATruseq_1ug |
| Stranded mRNA TruSeq | Illumina | mRNA | 100 | yes | ssmRNATruseq_100ng |
| Stranded total RNA TruSeq | Illumina | total RNA | 100 | yes | ssTotRNATruseq_100ng |
| **SMARTer technology** |  |  |  |  |  |
| RiboGone Stranded SMARTer | Takara Bio | total RNA | 100 | yes | RiboG_ssTotSmarter_100ng |
| RiboZero Gold stranded SMARTer | Illumina + Takara Bio | total RNA | 100 | yes | RiboZ_ssTotSmarter_100ng |
| RiboZero Gold stranded SMARTer | Illumina + Takara Bio | total RNA | 10 | yes | RiboZ_ssTotSmarter_10ng |
| **SMARTer Ultra-Low technology** |  |  |  |  |  |
| SMARTer UL + Nextera | Takara Bio + Illumina | mRNA | 1 (0.75) | no | mRNASmarterUL_XT1ng_750pg |
| SMARTer UL + Nextera | Takara Bio + Illumina | mRNA | 1 (0.13) | no | mRNASmarterUL_XT1ng_130pg |

### Reads quality metrics

Figure 1A shows the total number of reads mapped to the genome for the whole set of samples. All the 1 *μ*g samples (mRNA TruSeq, stranded mRNA TruSeq and stranded total RNA TruSeq categories) display close values (average values per class of 53, 49 and 55 M reads respectively). The 100 ng samples give average values similar to the 1 *μ*g samples (average values of 50, 52 and 52 M mapped reads for stranded mRNA TruSeq, stranded total RNA TruSeq and RiboGone stranded total RNA SMARTer categories), with the exception of the RiboZero stranded total RNA SMARTer samples, which display a much lower output at an average of 31 M reads. The 10 ng input category (RiboZero stranded total RNA SMARTer) shows the lowest value with an average of 22 M reads. The two ultra-low samples, mRNA SMARTer UL 750 pg and 130 pg yielded a relatively high number of reads with an average of 45 and 42 M reads.

**Figure 1.**
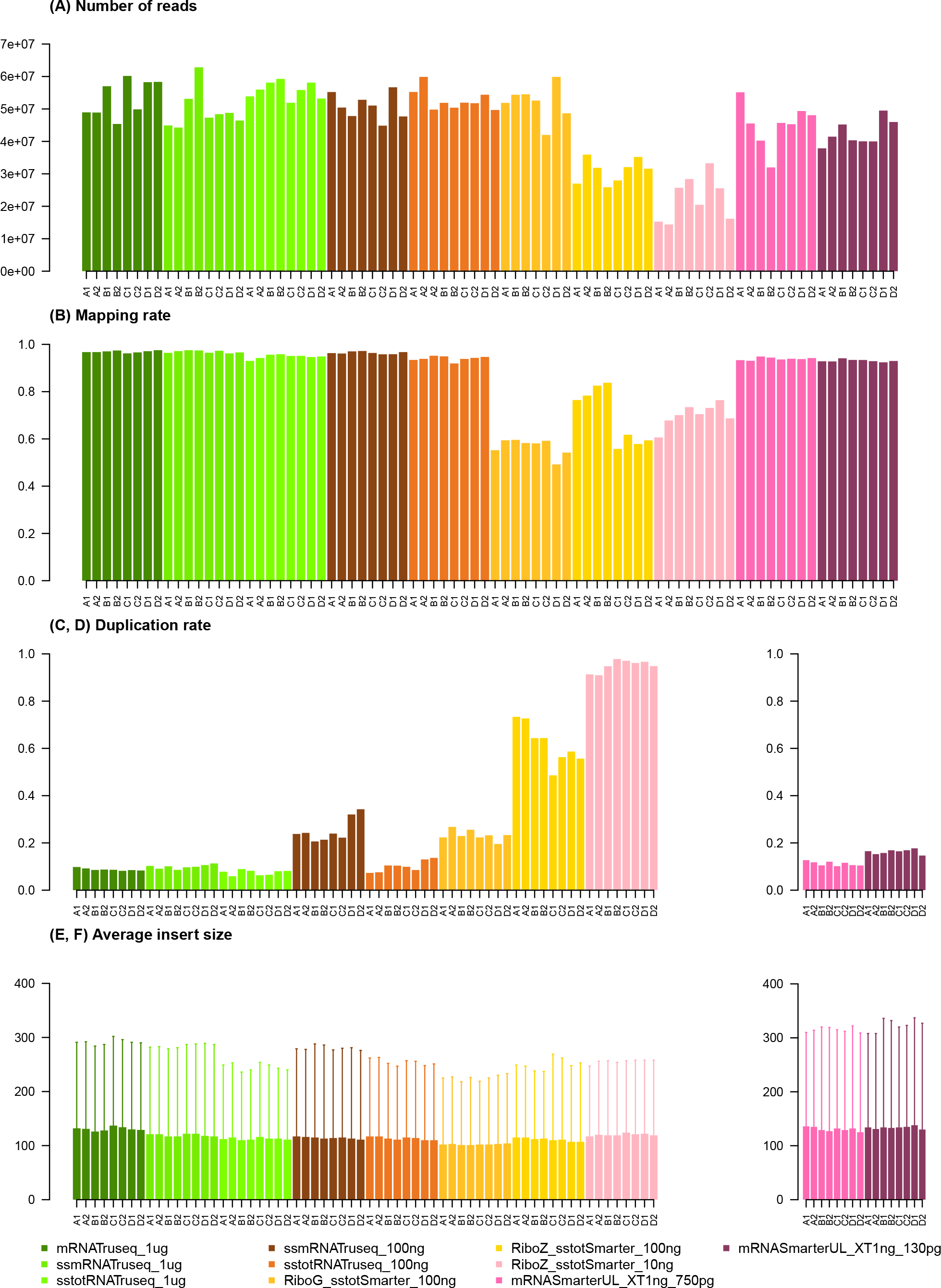
Reads quality metrics for all samples and conditions.

Figure 1B, shows the mapping rate for the different categories. Not surprisingly, the three 1 *μ*g categories have very high mapping rates with an average percentage of 95-96% of all reads mapped. For the 100 ng samples, two categories show a high percentage of read mapping (stranded mRNA and total RNA TruSeq), while the other two categories show lower mapping percentages (56% and 69% for the RiboGone and the RiboZero stranded total RNA SMARTer respectively). The 10 ng category shows an average mapping percentage of 69%, while the two SMARTer ultra-low samples (750 pg and 130 pg) display much higher mapping percentages at around 93%.

Duplication rates are shown in Figures 1C and 1D. Unsurprisingly, the rates are very low for all the 1 *μ*g samples (around 8% on average for all three categories). As expected the duplication rates for the 100 ng samples are higher, howerver large variations in average values can be seen depending on the kit used. The lowest average rate is 10% for the category stranded total RNA TruSeq, followed by stranded mRNA TruSeq and RiboGone stranded total RNA SMARTer at 25% and 23% respectively. The last 100 ng group displays a very high rate at 61% (RiboZero stranded total RNA SMARTer). The 10 ng samples have an extremely high rate of duplication with an average of 94%. The two ultra-low sample groups shown in Figure 1D have relatively low duplication rates with average values of 11% and 16% (750 pg and 130 pg respectively). However, caution should be taken in interpreting these values since the ultra-low kit has two consecutive steps of PCR amplification, and the values observed here represent duplication rates for the second stage only (see methods).

For the average insert size (Figure 1E and F), we observed very little variation between the different categories, with an overall average insert size of around 110 nt.

Globally, the results show that the 1 *μ*g groups display the best quality metrics results in terms of the high number of reads produced, as well as very high mapping rates and low duplication rates. No real distinction can be made at this level between the stranded versus unstranded or mRNA versus total RNA conditions. This group can be considered as the reference group in terms of performance and acceptable metrics levels. For the 100 ng input groups, the results are more contrasted. The two TruSeq groups (stranded mRNA and stranded total RNA) performed well, with a total number of reads and mapping rate comparable to that of the 1 *μ*g groups. Their duplication rates were higher than for the 1 *μ*g groups, although still in the acceptable ranges. The RiboGone stranded total RNA SMARTer group had a significantly lower mapping rate, while the last group, the RiboZero stranded total RNA SMARTer showed the lowest number of reads, a relatively low mapping rate and a very high duplication rate. Quality metrics for the 10 ng samples show very degraded results in all samples compared to the reference 1*μ*g group. The number of reads is extremely low, the mapping rate is relatively low and the duplication rate very high. On the other hand, the metrics for the two ultra-low input amounts were reasonably good compared to the reference samples, with a relatively high number of reads and a very high mapping rate and low duplication rate, although this last metric should be interpreted cautiously for this group, for the reasons mentioned above.

Figure 2 shows the normalized rates of mapping to intergenic, exonic and intronic regions of the genome. As could be expected, there is a clear distinction between the mRNA and total RNA samples. The mRNA samples all map predominantly to exonic regions, then to intronic and finally to intergenic regions. On the other hand, with the total RNA samples an almost equal proportion of reads map to the exonic and intronic regions, and considerably less reads map to the intergenic regions (although this smaller proportion is still higher than for mRNA samples). All the mRNA samples have similar proportions, with a slight decrease in the exonic regions for the ultra-low (SMARTer UL) groups. For the total RNA samples, the rates are very similar for the different kits and input amounts.

**Figure 2.**
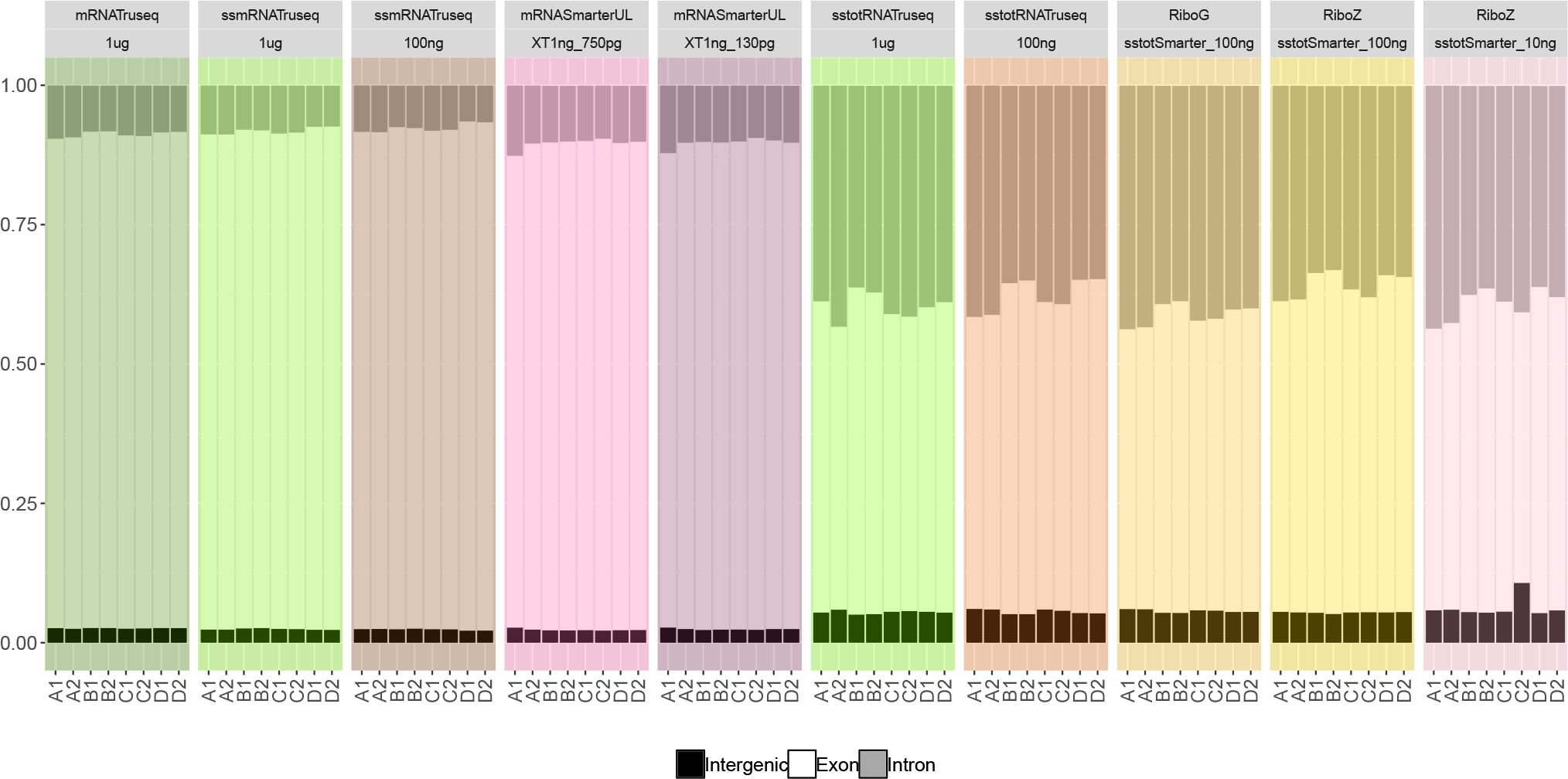
Normalized alignment rates to intergenic, exon and intron regions for all the samples. Note that the mRNA samples are on the left side while the total RNA samples are located on the right side of the figure.

### Gene detection and counting

Figure 3 shows the number of genes detected for each sequencing protocol and sequencing depth (2×2, 2×5, 2×10 and 2×25 M reads). We used a low threshold whereby a gene was considered expressed if it had a cpm (counts per million) value greater than or equal to one in two replicates (see methods). Overall, the different groups showed similar values for the number of genes detected. As also shown in Figure 3, the number of detected genes increases with the sequencing depth for all the categories, and the 2×2M sequencing depth category has the lowest number of detected genes in all the groups. Only the ultra-low input samples (mRNA SMARTer UL 750 and 130 pg) show a slight decrease in the number of genes detected. If we take the 1 *μ*g (stranded mRNA TruSeq and stranded total RNA TruSeq) and the 100 ng (stranded mRNA TruSeq, and stranded total RNA TruSeq) categories shown in figures 4A and 4B, we can see that the total RNA groups contain more detected genes than the mRNA samples, and that the proportion of non-coding RNAs and pseudogenes is higher for the total RNA groups. This is easily explained by the fact that non-coding transcripts are much better covered by total RNA protocols.

**Figure 3.**
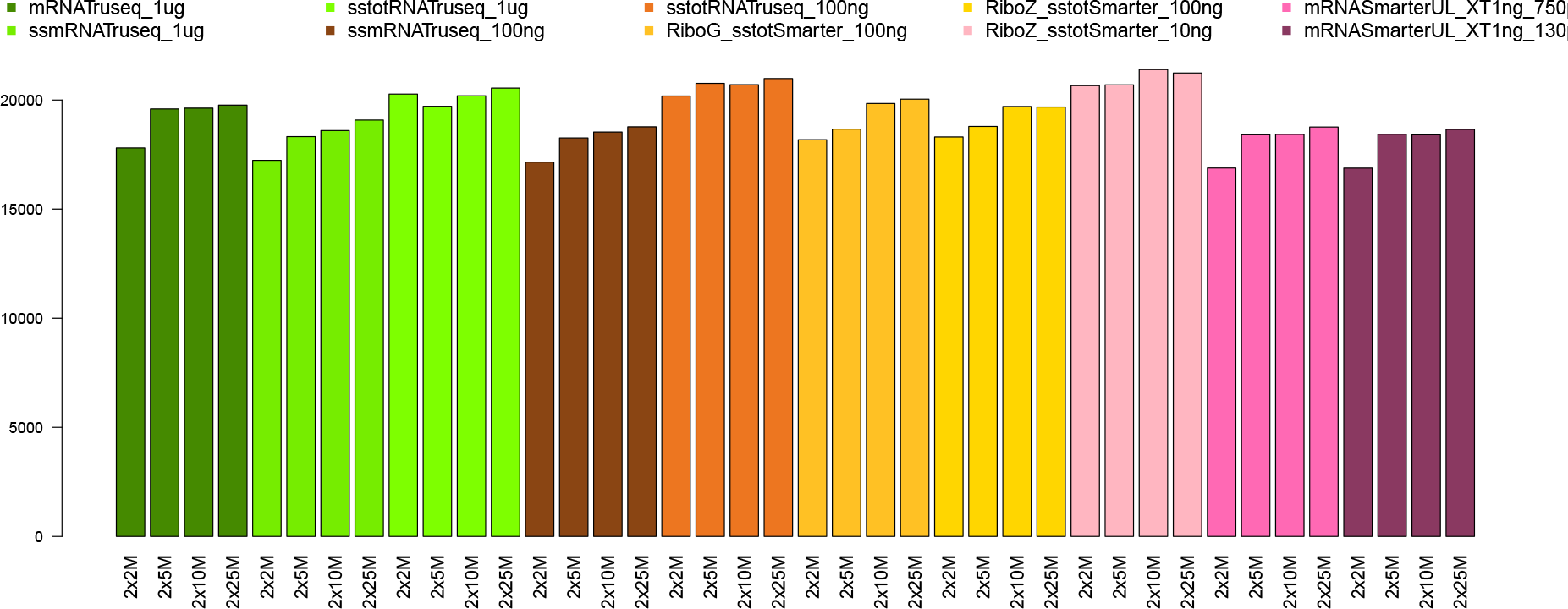
Number of detected genes for all the categories.

**Figure 4.**
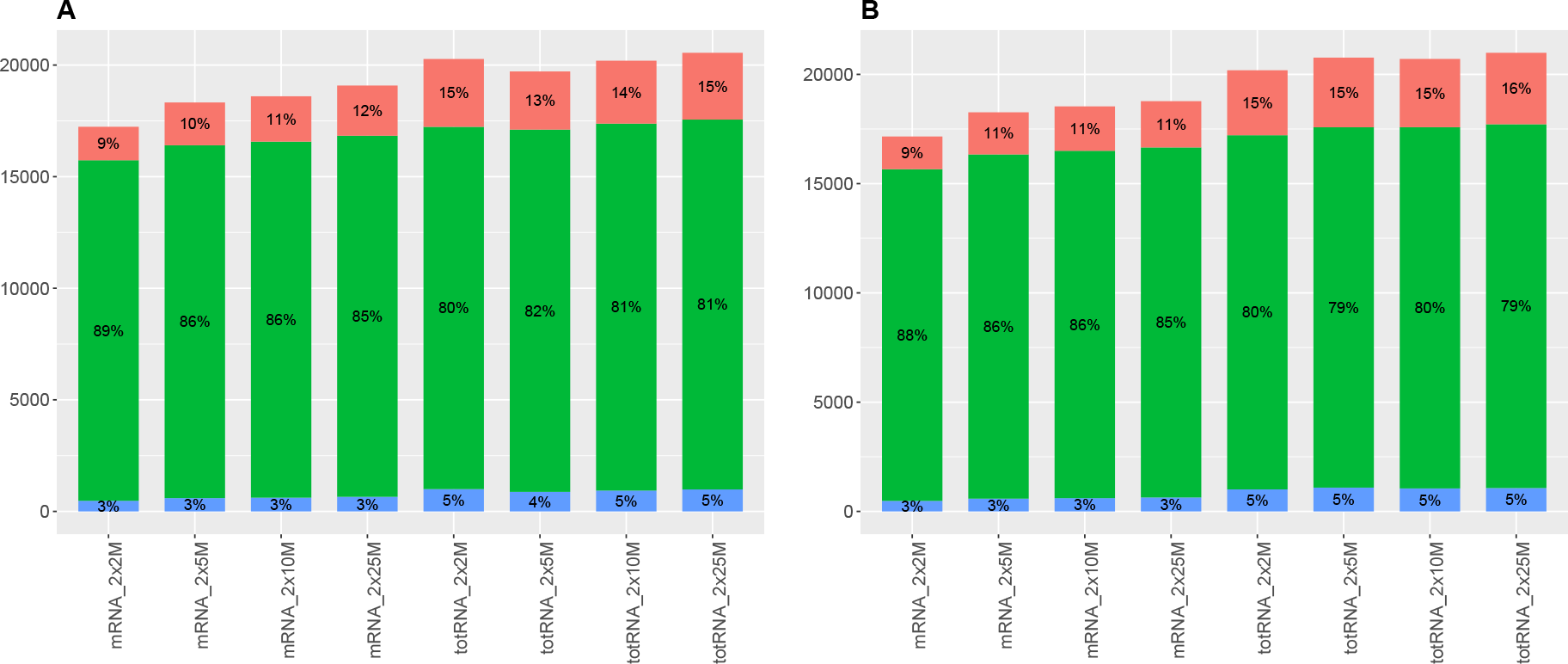
Percentage of pseudogenes (blue), protein-coding (green) and non-coding RNAs (red) for 1 *μ*g TruSeq samples (A) and 100 ng TruSeq samples (B). Note that mRNA samples are on the left side and total RNA samples are on the right side for each figure.

### Differential expression

Figure 5 shows the number of differentially expressed genes (DEG) in sample A (mixture of nomal human tissues) and B (human brain tissues) replicates. For the 1 *μ*g unstranded mRNA samples, a minimum of 11,059 DEG can be seen for the 2×2M sampling level and a maximum of 17,855 DEG for the 2×15M sampling level. The number of DEG is similar for the 1 *μ*g stranded mRNA, while a slight decrease can be observed for the 1 *μ*g total RNA samples: this is visible at all sampling levels. For the 100 ng samples, the DEG profiles observed for the stranded mRNA and total RNA TruSeq are very similar to those of their 1 *μ*g counterparts. In contrast, there is a clear decrease in the number of DEG detected for the 100 ng RiboGone and RiboZero stranded SMARTer samples, at all sampling levels. Out of all the groups, the 10 ng RiboZero stranded SMARTer samples clearly show the lowest number of DEG for all sampling levels. The number of DEG for the ultra-low samples is clearly higher at all sampling levels compared to the 10 ng RiboZero stranded total RNA SMARTer samples. The number of DEG in the ultra-low samples is comparable to that of the 100 ng RiboZero stranded total RNA SMARTer, with even higher levels observed for sampling levels 2×2M and 2×5M. Overall, despite their very low input amounts, the ultra-low samples performed reasonably well for the detection of DEG.

**Figure 5.**
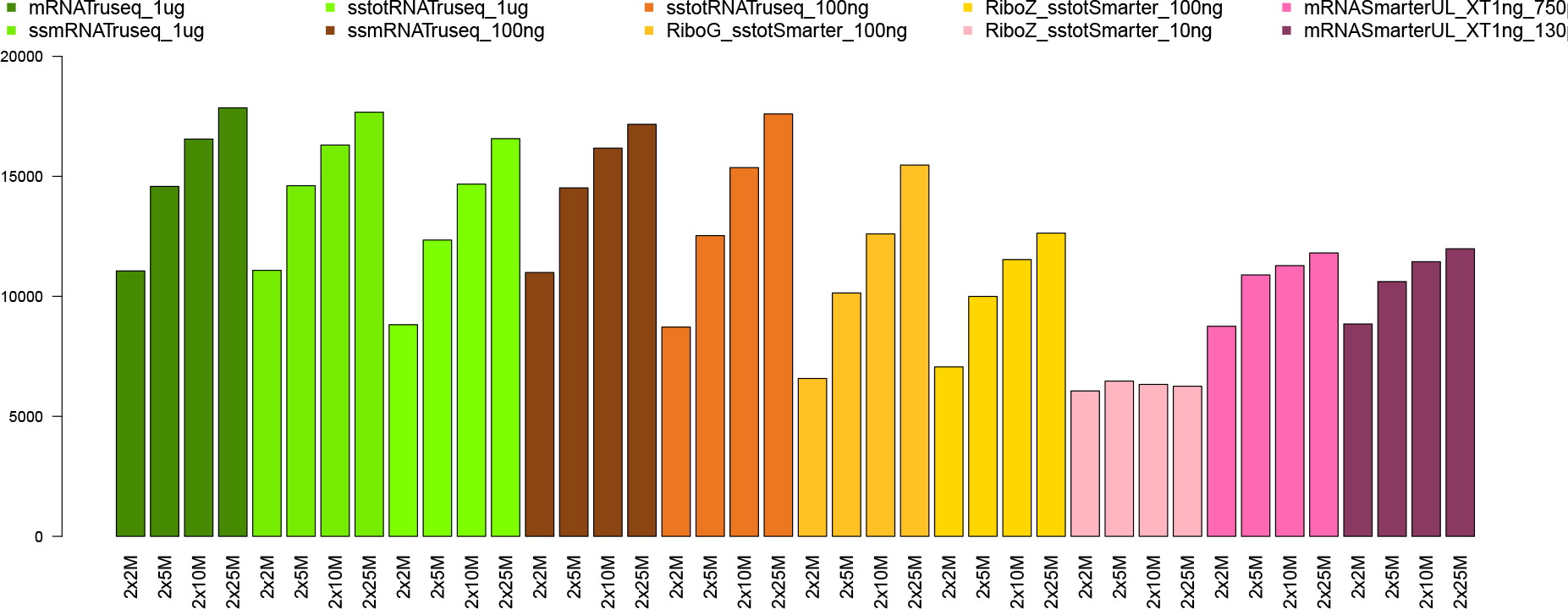
Number of differentially expressed genes (DEG) detected for all input quantities, preparation kits and sampling levels.

Table 2 shows an overlap in the number of DEG between the unstranded mRNA TruSeq 1 *μ*g samples used as the reference set and a selection of kits / input amounts. The overlap with the reference set is highest for the 1 *μ*g input amounts (91% and 79%), but is still relatively high for the 100 ng categories (89% and 84%), and, as expected, is lower for the ultra-low input quantities (54% and 55%), but still above 50%. This is not the case for the 10 ng RiboZero stranded total RNA SMARTer which falls below this level to 26%, thus much lower than the ultra-low samples.

**Table 2.**
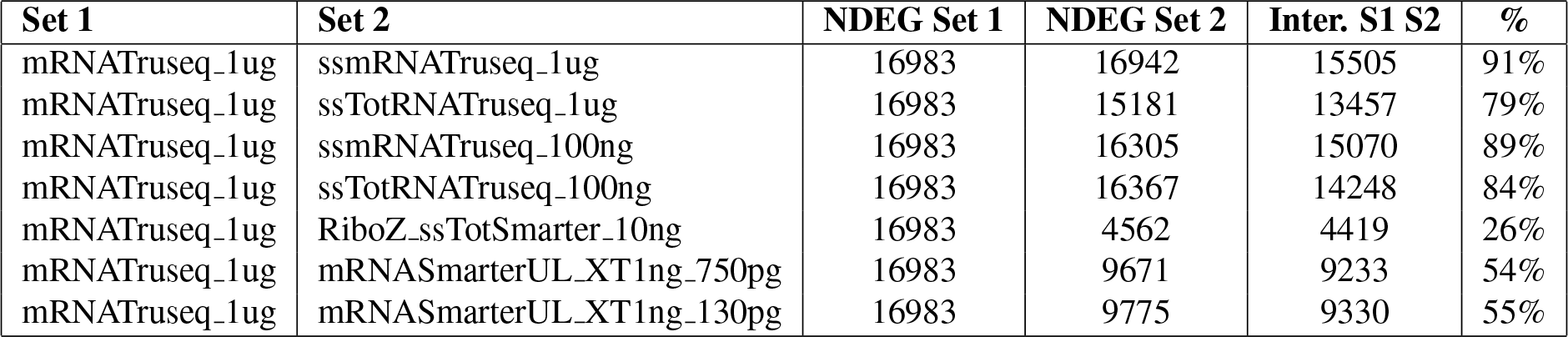
Overlap in the number of differentially expressed genes (DEG) between a reference set and a number of selected conditions for sampling level of 2×25M. The percentage represents the fraction of DEG relative to the reference set.

| Set 1 | Set 2 | NDEG Set 1 | NDEG Set 2 | Inter. S1 S2 | % |
| --- | --- | --- | --- | --- | --- |
| mRNAtruseq_1ug | ssmRNAtruseq_1ug | 16983 | 16942 | 15505 | 91% |
| mRNAtruseq_1ug | ssTotRNAtruseq_1ug | 16983 | 15181 | 13457 | 79% |
| mRNAtruseq_1ug | ssmRNAtruseq_100ng | 16983 | 16305 | 15070 | 89% |
| mRNAtruseq_1ug | ssTotRNAtruseq_100ng | 16983 | 16367 | 14248 | 84% |
| mRNAtruseq_1ug | RiboZ_ssTotSmarter_10ng | 16983 | 4562 | 4419 | 26% |
| mRNAtruseq_1ug | mRNASmarterUL_XT1ng_750pg | 16983 | 9671 | 9233 | 54% |
| mRNAtruseq_1ug | mRNASmarterUL_XT1ng_130pg | 16983 | 9775 | 9330 | 55% |

### Gene expression accuracy

The samples labelled C and D are a mixture of the Human Universal Reference Total RNA (A) and Human Brain Reference RNA (B), with ratios of 75% A + 25% B and 25% A + 75% B respectively. Since we have the gene expression levels for samples A and B, we can predict the theoretical gene expression values for the mixture samples C and D and compare these to the values obtained by analyzing the samples. Thus we can evaluate the accuracy of the gene expression levels in samples C and D for all the conditions tested. The boxplots in figures 6 and 7 show the gene expression levels ratios between the real and predicted values for samples C and D respectively. The red dotted line in the figures indicates the perfect agreement (ratio value of 1) between the real and the predicted values. For the sample C ratios (Figure 6), in most cases the median value for each group is aligned or very close to the red dotted line, indicating very good agreement between the predicted and real gene expression values. For the sample D ratios, the agreement is globally good, but as can be seen from the figure there are larger deviations from the red dotted line, for instance for the TruSeq stranded total RNA 1 *μ*g group, the TruSeq stranded mRNA 100 ng group, the RiboZero stranded total RNA SMARTer 10ng group and the mRNA SMARTer UL 130 pg. We also clustered samples A, B, C and D according to the normalized counts and in most cases, the samples clustered logically, showing the C samples to be more closely related to A samples, and the D samples more closely related to the B samples (see supplementary figure 1 for two selected examples).

## Discussion

Following global cost reductions for sequencing, RNA-seq is now an affordable and popular technique for various areas of biological research, with potential applications in clinical medicine^1^. For instance, RNA-seq can be applied in clinical context for the detection of aberrant transcription in human diseases such as aberrant expression profiles, gene fusion expression, alternative transcripts and allele-specific expression^1^. However, robust gene expression analysis and other potential clinical applications of RNA-seq are dependent on the quality, reliability and reproducibility of the extraction and sequencing procedures. In this study, we performed a comprehensive assessment of several state-of-the-art and popular commercial kits for RNA-seq library preparation kits using standard, low and ultra-low input amounts. We assessed the performance of the kits using various sequencing and alignment read metrics, gene expression detection levels and gene differential expression levels at various sampling depths. The results are summarized in table 3, which indicates the global levels of quality for the different metrics analyzed in this study. We also included an indicator of the automation capability of each kit, i.e. to indicate whether or not the kit can be used with liquid handling robotic systems. This capacity can be important for the reproducibility and robustness of the results, in the case of large research projects or in sequencing centers that process large numbers of samples.

**Table 3.**
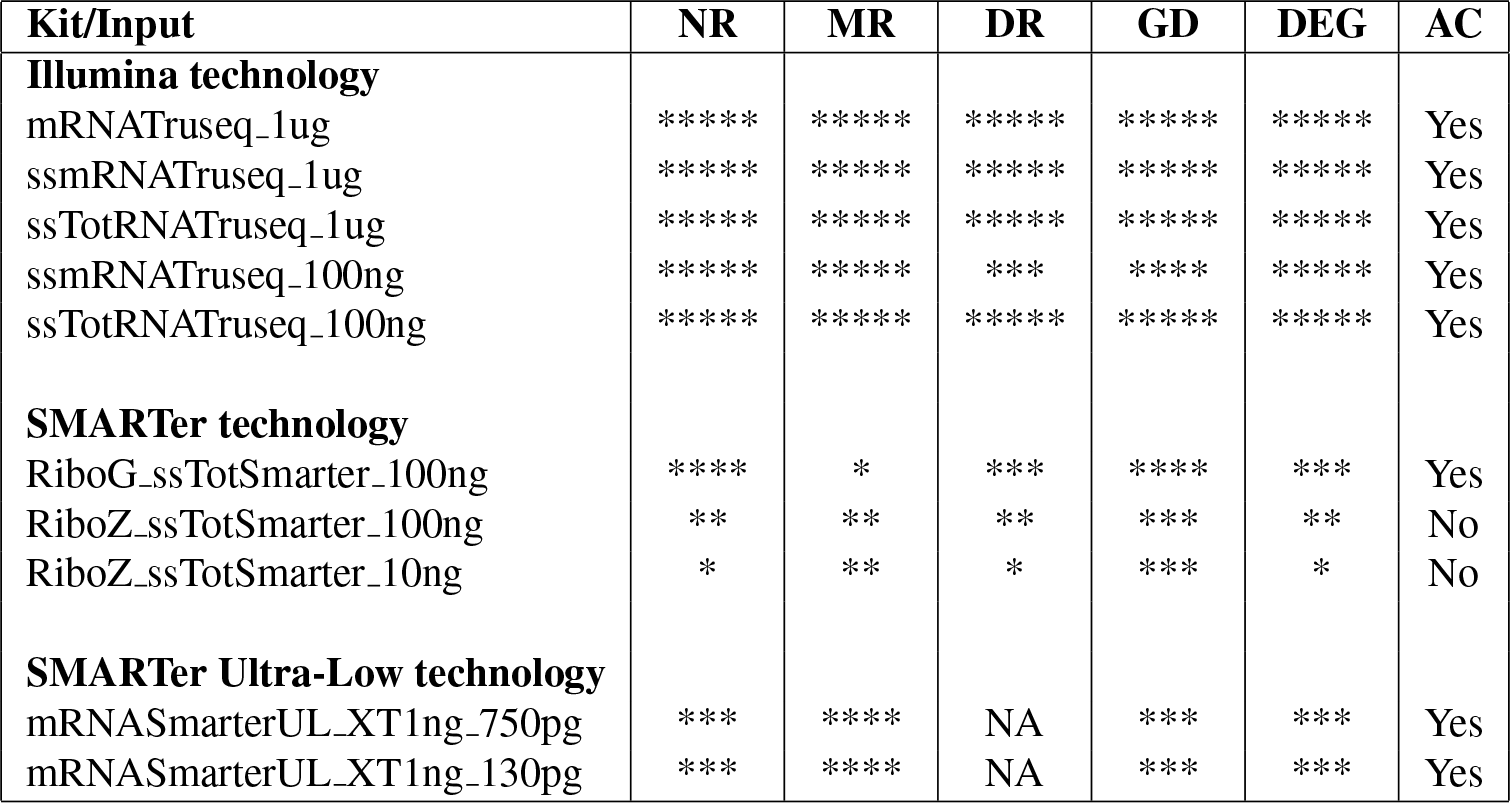
Assessment summary of the different RNA preparation kits and conditions analyzed in this study. For clarity, the table presented here is based on the 2×10M sampling level. The number of stars indicates the global level of quality for the metrics used, with a maximum of five stars for best quality. NR, MR and DR: sequencing and alignment quality metrics (Number of reads, mapping rate and duplication rate). GD: gene detection metrics. DEG: differentially expressed gene metrics. AC: automation capability.

**Figure 6.**
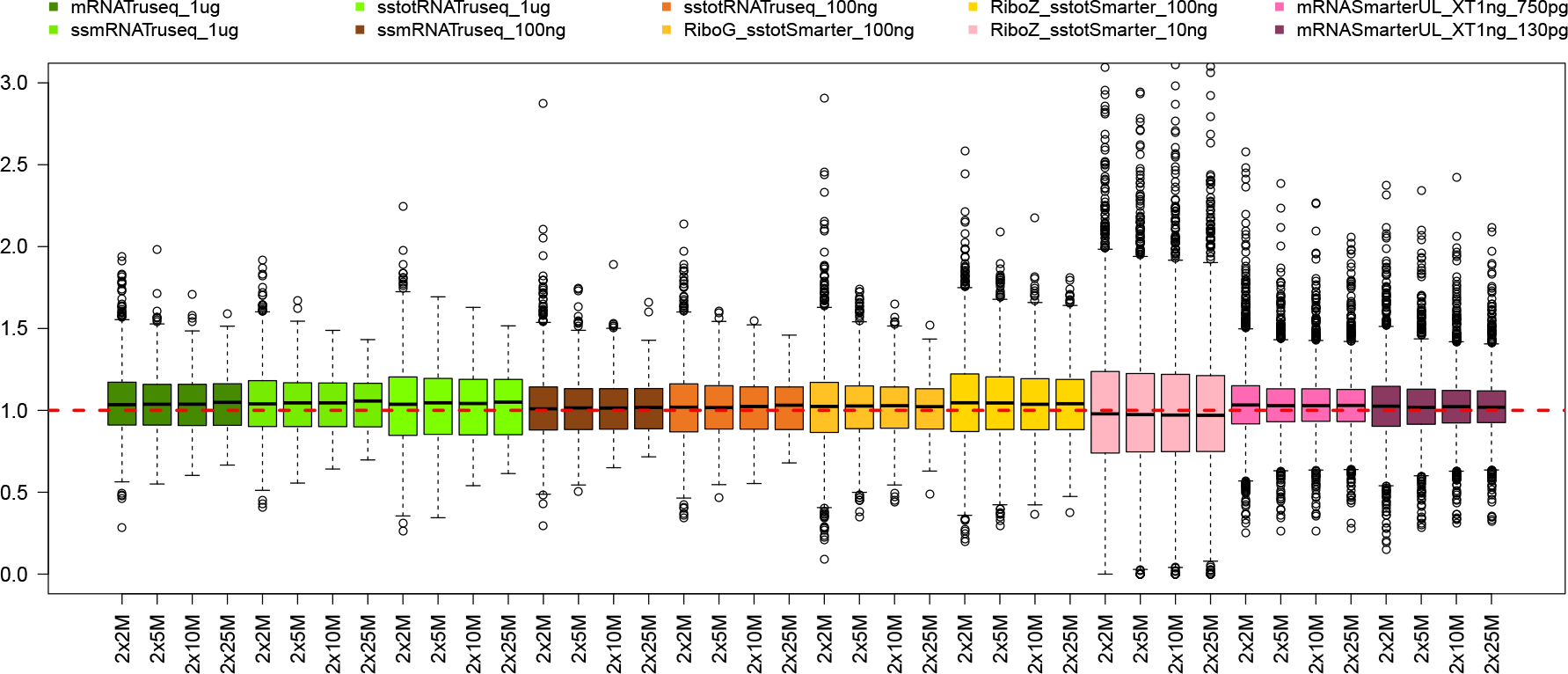
Ratios between real and predicted gene expression values for samples C for all groups. The red line indicate a ratio value of 1, i.e. a perfect match between real and predicted value. Some of the outlier values are not shown on this figure.

**Figure 7.**
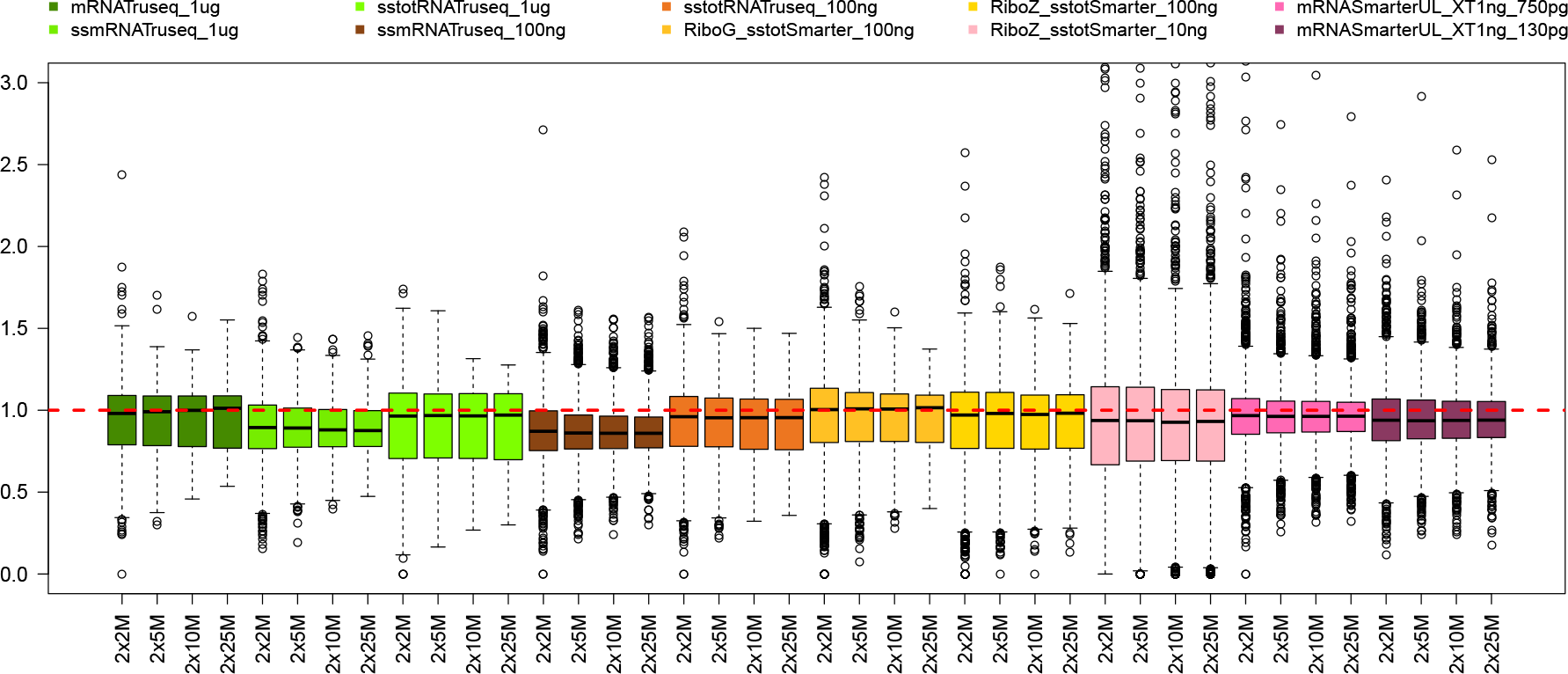
Ratios between real and predicted gene expression values for samples D for all groups. The red line indicate a ratio value of 1, i.e. a perfect match between real and predicted value. Some of the outlier values are not shown on this figure.

## Conclusion

The TruSeq unstranded mRNA 1 *μ*g kit can be considered as the reference material or the “gold standard” in this study, as this kit has been very widely tested and used for RNA-seq studies, and the 1 *μ*g amount of input material is usually considered more than enough for most RNA-seq applications. The other kits used for the 1*μ*g input amount (stranded TruSeq mRNA and total RNA) perform equally well in terms of the sequencing and alignment metrics, the number of detected genes and the number of genes differentially expressed. For the 100 ng input amounts, the TruSeq kits (stranded mRNA and total RNA) achieve similar performances, except for a slightly elevated duplication rate for the TruSeq stranded mRNA. The other kits tested for 100 ng input, i.e. the RiboZero or RiboGone stranded total RNA associated with the SMARTer kit, clearly show degraded performances on multiple levels compared to the TruSeq kits. For the low and ultra-low input amounts, the stranded RiboZero SMARTer kit with the 10 ng input amount underperforms on most of the indicators. In contrast, the unstranded mRNA SMARTer samples (750 pg and 130 pg), with single cell-like input levels quantities, achieve very good levels for most of the performance indicators, and particularly for the sequencing and alignment quality metrics.

## Methods

### Samples

Two commercial RNA samples were used to evaluate the different RNA-seq applications. Human Universal Reference Total RNA (Takara Bio/Clontech, labeled A), a mixture of 23 normal human tissues (including brain) and First Choice Human Brain reference RNA (Ambion), a pool of normal human brain tissues (labeled B). The other samples were prepared as follows: C = 75% A + 25% B; D = 25% A + 75% B. All the samples were prepared in duplicate for all the RNA applications.

### RNA-seq protocols

#### Illumina TruSeq technology

Three kits from Illumina were tested with different total RNA input: TruSeq mRNA with an input amount of 1 *μ*g, TruSeq stranded mRNA with inputs of 1 *μ*g and 100 ng and TruSeq stranded total RNA Gold with 1 *μ*g and 100 ng. The TruSeq mRNA and stranded mRNA kits use polyA selection to trap the mRNA before proceeding to the cDNA synthesis. In the case of the TruSeq stranded total RNA Gold kit, the RiboZero Gold technology targets the rRNA with baits, and the cDNA synthesis takes place after purification. To ensure that the information on the strand is kept in the experimental procedure, the stranded kits use both actynomycin-D (during the first strand cDNA synthesis) and dUTP (during the second strand synthesis). The libraries are end-repaired and adenylated before being ligated with Y-shape single indexed adaptors, and then are amplified by PCR (Supplementary Table S1). We followed the Illumina recommendations for all the kits, except for the last purification step which we performed with Ampure XP beads at 0.8X to remove all the adapter dimers.

#### Stranded SMARTer technology

Starting with mRNA or ribosomal depleted RNA material, the stranded SMARTer technology (Takara Bio/Clontech) is based on tagged random hexamers primers and a SMARTScribe Reverse Transcriptase (RT) with terminal transferase activity. When it finishes the first strand cDNA synthesis, the RT adds a few non-templated nucleotides to the 3’ end of the cDNA (GGG). The SMART adapter is complementary to these nucleotides and adds the 5’ tag. The first strand cDNA is used as a matrix to perform the PCR amplification using indexed primers. In order to remove the rRNA from the input, we tested the RiboZero Gold kit (Illumina) and the recently released RiboGone Mammalian kit (Takara Bio/Clontech). The latter combines hybridization of rRNA and mtRNA baits with RNAse H digestion. A combination of RiboZero plus SMARTer stranded was used with 10 ng and 100 ng of total RNA input. Purification was performed on purifying columns (Macherey Nagel) before using the SMARTer stranded kit. We also tested a combination of RiboGone plus SMARTer stranded with an input of only 100 ng. We followed the Takara Bio/Clontech recommendations and the number of PCR cycles that we performed is indicated in Supplementary Table S1.

#### SMARTer Ultra Low technology

The SMARTer Ultra Low technology (Takara Bio/Clontech) is also based on the RT properties but uses polydT primers to synthesize the first strand of the cDNA. The full-length cDNA is then amplified by LD-PCR before being quantified with the High Sensitivity Qbit kit and qualified with the High Sensitivity chip in a BioAnalyzer machine. We normalized in 2 different quantities: 130 pg and 750 pg were used as input for the Nextera XT kit (Illumina). The Nextera technology uses a modified tranposase to fragment and ligate adapters in a unique step called “tagmentation”. The libraries are then amplified by PCR using indexed primers. Full details of the LD-PCR and PCR are given in Supplementary Table S1. The last purification step is performed with a ratio of 0.6X of XP Ampure Beads.

#### Sequencing

The qualification and quantification estimations for each library were done after the last purification using DNA1000 chip (input ≥ 100 ng) or High Sensitivity chip (input < 100 ng) for BioAnalyzer (Agilent). After normalization the libraries were sequenced in 4-plex on HiSeq 2000 or HiSeq 2500 machines (Illumina) in paired-end 2 × 100 bp.

### Quality check, alignment and quantification

After sequencing, read quality was checked with fastQC^21^ (version 0.11.3) and trimmed with a quality threshold of 30 using trimmomatic^22^ (version 0.32). The reads were then aligned against the Ensembl human reference genome GRCh37 with Tophat2^23^ (version 2.0.13). We used the Picard-Tools^24^ (version 1.138) module CollectRNASeqMetrics to calculate the mapping rate, the duplication rate and the average insert size. Finally, we used htseq-count^25^ (version 0.6.1) to assign and count the reads for all the Ensembl GRCh37 gene models.

#### Gene expression detection

Expression detection sensitivity was based on the analysis of the two technical A replicates for each condition. All genes for which both cpm (counts per million) of normalized expression were greater than 1 were considered as expressed. In other words, we declared as detected all genes for which:

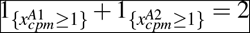

#### Genes differential expression

All statistical analyses were performed with the R software^26^ (version 3.4.3). Differential expression between the two HsRef replicates (A) and the two Brain replicates (B) was studied using the R package EdgeR which implements a TMM normalization and a negative binomial approach as described by Robinson et al.^27^. We removed from the analysis all genes for which all cpm values (for the four samples) were lower than 1. To take into account the multiple testing problem that arises when testing a large number of genes simultaneously, we applied the Benjamini-Hochberg procedure (BH) at a 5% threshold.

It is worth noting that we also considered the two other popular approaches for differential analyses, DESeq2^28^ and voom+Limma^29^, that lead to similar results.

#### Gene expression accuracy

To evaluate the accuracy of each protocol, we considered only genes which were declared as differentially expressed for all protocols in the previous step. To take account of differences in gene length, we normalized the raw counts using the FPKM approach. Predicted values (*Ĉ* and *D̂*) were calculated as the weighted sum of the averages of the two replicates for A and for B:

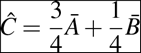

and

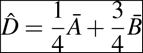

For each gene, we then calculated the difference between the mean (over the two replicates) of the observed and predicted values for the mixture C (resp. D) and the predicted values *Ĉ* (resp. *D̂*). We also calculated both the euclidean distance and the Pearson correlation coefficient between the average normalized measurements for C (resp. D) and the predicted values *Ĉ* (resp. *D̂*). Finally, we performed hierarchical clustering using all genes with Euclidean distance and average linkage, to determine whether similar samples clustered together.

## Acknowledgements

We would like to thank Elizabeth May for proofreading the manuscript.

## Author contributions statement

R.O., C.D. and MA.P. designed the study. MA.P and C.D. performed the experiments. MA.P., Cy.D., E.B., C.B. and S.BJ. wrote the scripts and analyzed the data. R.O., JF.D., C.A. discussed the results and commented on the manuscript. E.B., MA.P. and Cy.D. wrote the manuscript. All authors reviewed and approved the manuscript.

## Additional information

**Supplementary Table 1:** number of PCR cycles done for each application. For the Ultra Low Smarter v4 followed by Nextera XT, the first number is for the LD-PCR while the second number is for the PCR.

**Supplementary Figure 1:** hierarchical clustering of samples A, B, C and D from gene counting data for the conditions unstranded mRNA Truseq 1 ug 2×25 M (A) and unstranded mRNA Smarter 1 ng (750 pg) 2×25M (B).

