## Supplementary table S1 for "Comprehensive analysis of RNA-seq kits for standard, low and ultra-low quantity samples"

Number of PCR cycles done for each application. For the Ultra Low Smarter v4 followed by Nextera XT, the first number is for the LD-PCR while the second number is for the PCR.

| **Applications** | **Input Total RNA** | **PCR cycles** |
|---|---|---|
| mRNA Truseq (Illumina) | 1000ng | 13 |
| Stranded mRNA Truseq (Illumina) | 1000ng | 13 |
| | 100ng | 15 |
| Stranded total RNA Truseq (Illumina) | 1000ng | 14 |
| | 100ng | 15 |
| RiboGone + stranded Smarter (Clontech) | 100ng | 18 |
| RiboZero Gold (Illumina) + stranded Smarter (Clontech) | 100ng | 18 |
| | 10ng | 18 |
| Ultra Low Smarter v4 (Clontech) + Nextera XT (Illumina) | 1ng/130pg | 11/12 |
| | 1ng/750pg | |

Supplementary Table 1 :
