## Supplementary figure 1 for "Comprehensive analysis of RNA-seq kits for standard, low and ultra-low quantity samples"

**A**

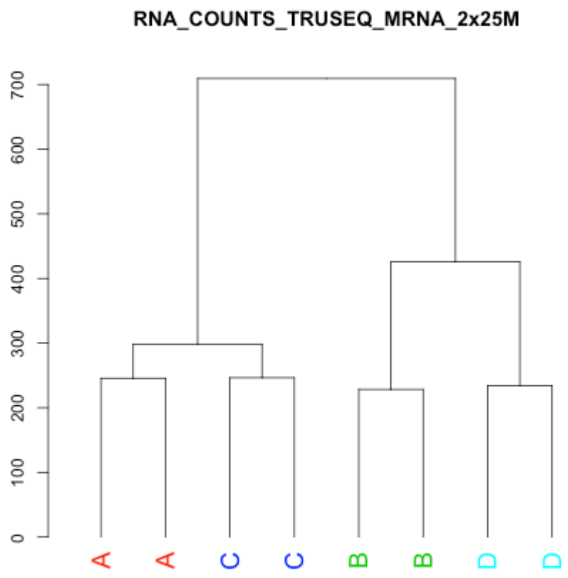


**B**

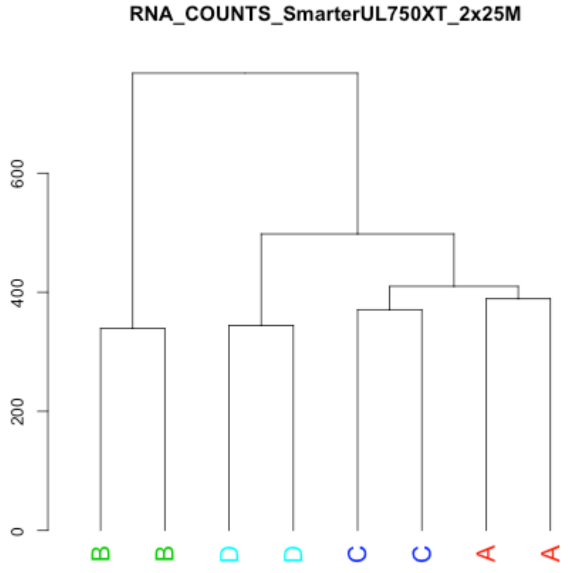


**Supplementary figure 1**: hierarchical clustering of samples A, B, C and D from gene counting data for the conditions unstranded mRNA Truseq 1 ug 2x25 M (**A**) and unstranded mRNA Smarter 1 ng (750 pg) 2x25M (**B**).
